# SunRISE: long-term imaging of individual mRNA molecules in living cells

**DOI:** 10.1101/2020.10.05.326660

**Authors:** Yue Guo, Robin E. C. Lee

## Abstract

Single-cell imaging of individual mRNAs has revealed core mechanisms of the central dogma. However, most approaches require cell fixation or have limited sensitivity for live-cell applications. Here, we describe SunRISE (SunTag-based Reporter for Imaging Signal Enriched mRNA), a computationally and experimentally optimized approach for unambiguous single-mRNA detection in living cells. We demonstrate SunRISE with long-term epifluorescence imaging, using translational stress to track mRNA phase separation and recovery from cytosolic droplets.

## Main

Messenger RNA (mRNA) molecules interact with RNA binding proteins throughout their lifespan to carry genetic information and provide precise spatiotemporal regulation within cells. Live-cell single molecule imaging techniques have enabled in-depth characterization of dynamics for mRNA processing steps, including transcription, translation, and interactions with ribonucleoprotein (RNP) granules^1-5^. However, continuous imaging of single mRNAs has numerous challenges coupled to low imaging sensitivity that is exacerbated by rapid photobleaching^6^. Typically, live-cell detection of single mRNAs requires sophisticated imaging approaches and trade-offs that restrict spatial and temporal aspects of imaging experiments.

Bacteriophage-derived MS2 and PP7 stem-loops are extensively used for labeling mRNA molecules. In many applications, the reporter mRNA is tagged with 24x stem-loop copies in the 3’ UTR and the corresponding coat protein (MCP or PCP) is fused with a fluorescent protein (FP). When co-expressed in the same cell, dimers of FP-fused coat proteins bind to each stem loop enabling visualization of mRNAs and active transcription sites by fluorescence microscopy^7^. Although variants with increased stem-loop numbers (MS2×128) enhance imaging sensitivity^8^, the bulky mRNA extension can significantly alter mRNA dynamics and the reporter still suffers from photobleaching.

We set out to develop a signal enrichment approach to circumvent these problems and facilitate long-term imaging of mRNAs in live cells. To amplify fluorescence intensity of labeled coat proteins, our design employs SunTag, an array of GCN4 peptide epitopes that recruit multiple antibody molecules^9^. Specifically, SunRISE (SunTag-based Reporter for Imaging Signal Enriched mRNA) comprises two stages of signal amplification: 24x stem-loop copies are inserted in the 3’-UTR of mRNA and the corresponding coat protein is fused with up to 24x SunTag peptides. With co-expression of an FP-fused single-chain antibody (scFv-GFP or codon optimized scAB-GFP^10^) that bind GCN4 epitopes, each SunTag-coat protein can be labeled with up to 24 scFv-FPs resulting in an upper bound of over 1000 FP molecules per mRNA (Fig. 1a). We reasoned that a two-stage approach would bolster active cycling of nascent coat proteins and FPs, thereby providing resistance to photobleaching while minimizing alterations to target mRNA.

**Fig. 1:**
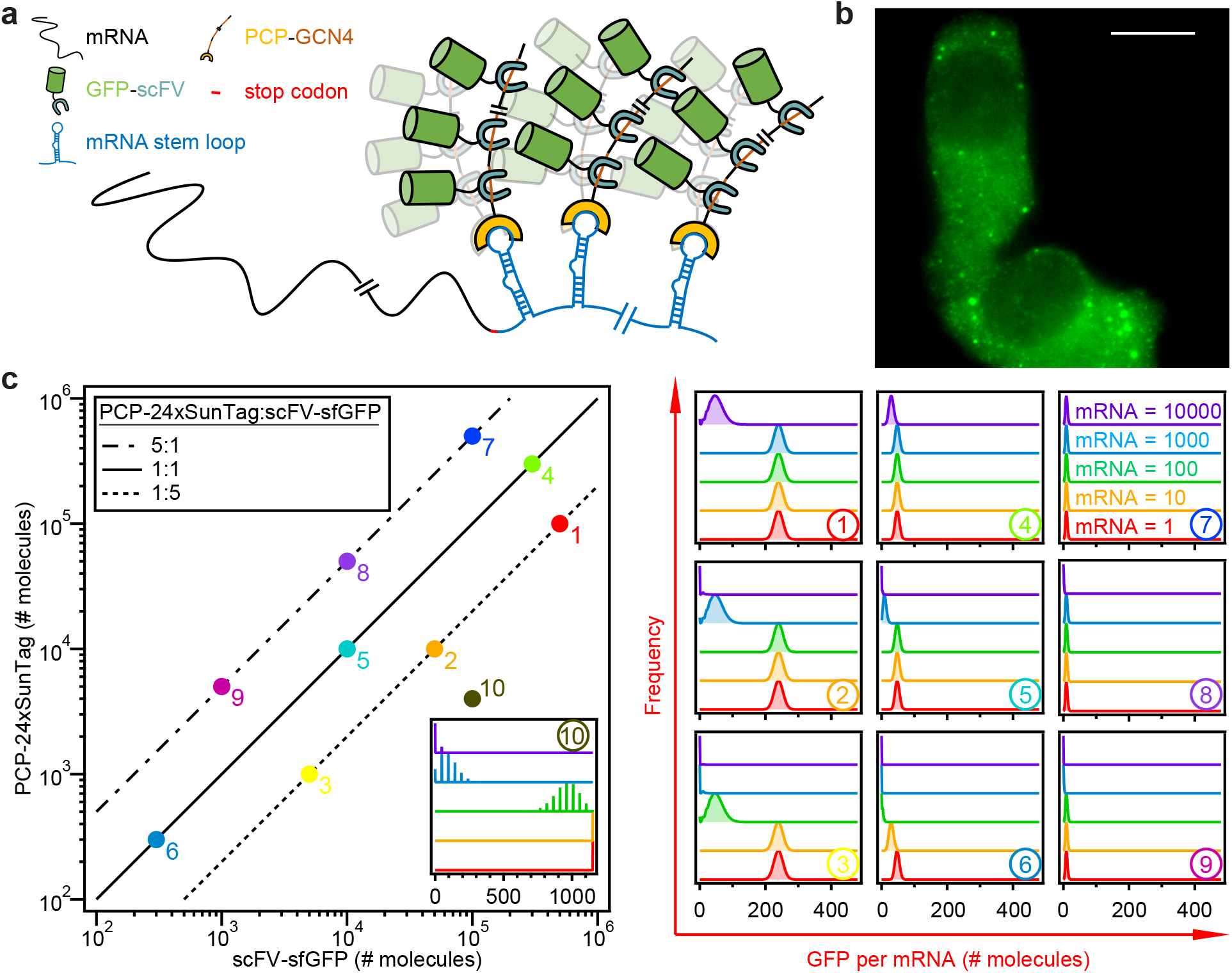
Design of SunRISE, a SunTag based reporter for imaging signal-enriched mRNA. **a**, Schematic of the SunRISE strategy for imaging single mRNA molecules. An mRNA transcript (black) tagged at 3’ UTR with PP7 stem-loops (blue) is bound by the PCP coat protein (yellow), which is fused to a SunTag GCN4 peptide array (orange). SunTag recruits GFP (green) through antibody-peptide specific binding between scFv (grey) and GCN4 epitopes. **b**, Maximum intensity projection of z-stacks for representative HeLa cells 24 hours after transfection with three plasmids: phage-cmv-cfp-24xpp7, cmv-24xSunTag-PCP and cmv-scAB-sfGFP. Scale bar: 10 μm. **c**, Parameter sweeps in the space spanned by number of molecules for scFv-GFP and 24xSunTag-PCP (left) using a computational model to calculate number of GFP molecules per mRNA. Parameter combinations 1-10 were selected to represent different ratios between scFv-GFP and 24xSunTag-PCP (solid line 1:1, dotted line 5:1 and dash-dotted line 1:5) and different expression levels and frequency plots are shown (right). Different concentrations of mRNAs per cell are distinguished by different colors.

We expressed components of the naive design in HeLa cells under control of the cmv promoter and imaged cells by 3-D epifluorescence microscopy. Although diffraction-limited spots were visible, we observed spot-to-spot variability in size and intensity (Fig. 1b). It was anticipated that two-stage signal amplification has non-linear systems properties, e.g. limiting scFv-GFP or excessive SunTag-PCP concentrations will both influence signal intensity dependent on mRNA concentration. We therefore developed a mathematical model (see Methods) to estimate the number of FP molecules bound to mRNA for different expression levels of SunRISE components, scAB-GFP, SunTag-PCP, and PP7-tagged mRNA. Using protein-binding kinetics for SunRISE components and sampling different parameter combinations, we found broad variability in the expected intensity and signal-to-background (Fig. 1c and Supplementary Fig. 1), in some cases leading to a quantized distribution of single mRNA intensities (e.g. parameter combination 10). Variability between spots will complicate accurate identification and measurement of single mRNA molecules. By inspection, we found that a 5:1 ratio between scAB-GFP and SunTag-PCP with high-abundance expression yields uniform signal intensity distributions in cells expressing up to 1000’s of mRNAs (Fig. 1c). We also simulated 5x, 10x and 24x SunTag-PCP variants. Although longer variants produce more intense signals, all had comparable signal-to-background ratios (Supplementary Fig. 1).

Guided by simulations, we designed SunRISE variants and assayed quantitative properties of mRNA spots. Comparing promoters in HeLa cells, we found that GFP expression from the cmv promoter is approximately 5 times that of ubc (Supplementary Fig. 2a). To reduce the size of labeled mRNA complexes without significantly compromising signal to background, we validated model predictions using 10xSunTag-PCP. HeLa cells were co-transfected with SunRISE components, approximating parameter combinations 1, 4, 6 and 9 (Fig. 1 and Supplementary Fig. 1). Consistent with simulations, an approximately 5:1 ratio achieved by cmv-scAB-GFP and ubc-SunTag-PCP enhanced signal intensity and signal-to-background values (Supplementary Fig. 2b). Using smFISH against PP7 stem-loop sequences^11^ in SunRISE-expressing cells, we confirmed co-localization of cytoplasmic mRNAs between SunRISE and smFISH. However, smFISH revealed nuclear mRNAs that were not detected by SunRISE (Supplementary Fig. 3a). Further live-cell and fixed-cell assays demonstrated that GFP-SunTag-PCP is excluded from the nuclear compartment (Supplementary Fig. 3), suggesting that SunTag-PCP required further optimization for whole-cell mRNA detection.

To alleviate nuclear export effects from repeats of the GCN4 epitope, we continued with the smaller 5xSunTag variant. Next, we focused on modifications to 5xSunTag-PCP to shift the balance between nuclear and cytoplasmic expression. We also considered that ornithine decarboxylase (ODC) tag^11^ fused to SunTag-PCP while under control of the cmv promoter as an alternative approach to maintain a 5:1 deficit of SunTag-PCP that may also alter its sub-cellular distribution. Although several variants enable whole-cell mRNA detection, an optimized design was eventually achieved by switching the fusion order of PCP and 5xSunTag, in addition to inserting a 5’ nuclear localization signal (NLS; Supplementary Fig. 4 and Supplementary Table 1). The optimized SunRISE design faithfully labels single mRNAs in the nucleus and cytoplasm with uniform fluorescence intensity and high signal-to-background, allowing for long term imaging (Fig. 2, Supplementary Fig. 5, and Supplementary Movie 1). Furthermore, the same optimization can be applied to orthogonal stem-loops and antibody-epitope pairs, such as MS2, MS2V6^12^, and MoonTag^13^ (Fig. 2, Supplementary Figs. 6 & 7).

**Fig. 2:**
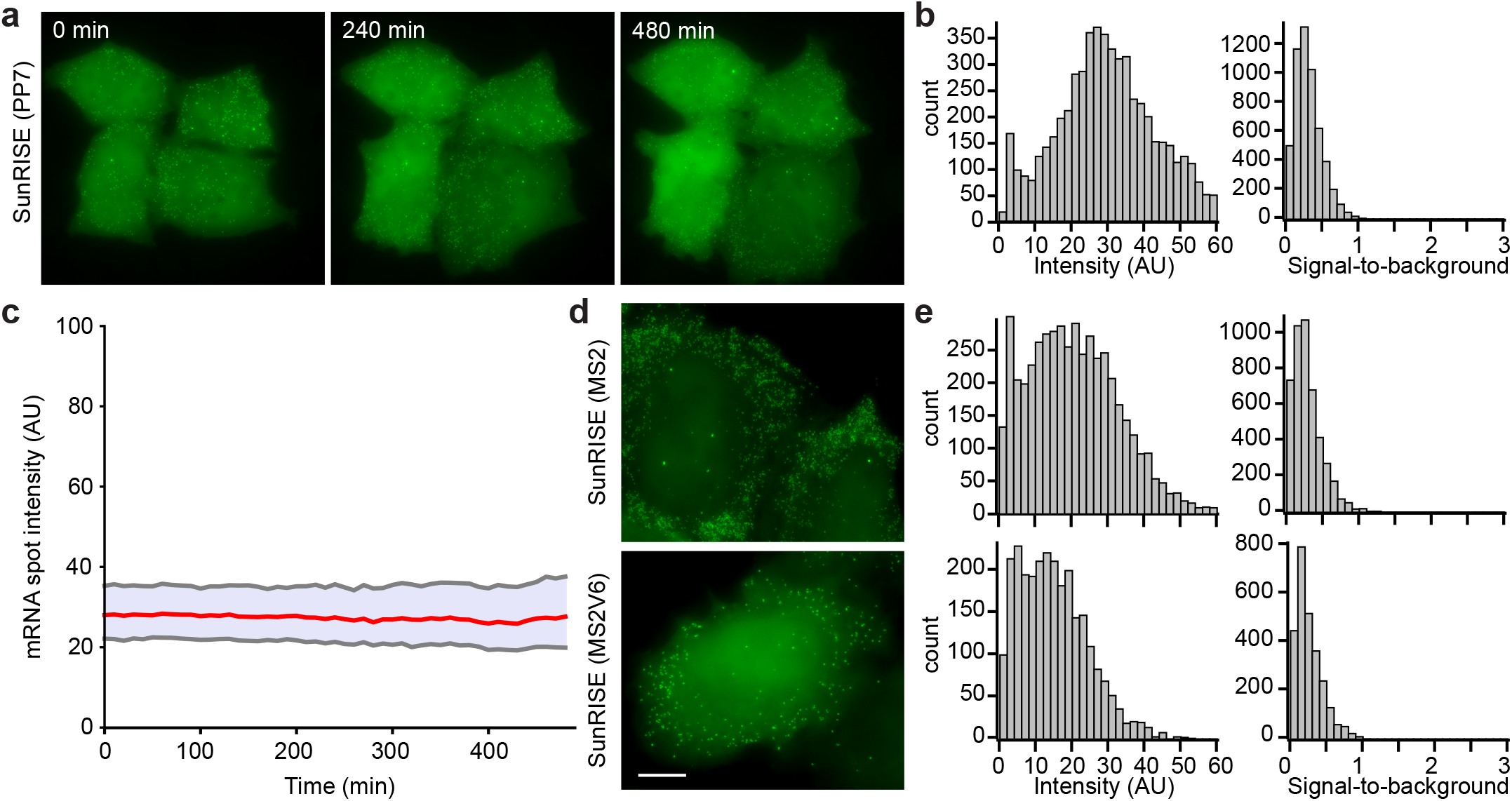
The optimal SunRISE design allows long-term imaging of single mRNA molecules throughout the cell. **a**, Maximum intensity projections of HeLa cells transfected with phage-cmv-cfp-24xpp7, ubc-nls-PCP-5xSunTag and cmv-scAB-sfGFP (SunRISE). Cells were imaged by 60x widefield fluorescence microscopy for 8 hours with a 10-minute framerate. Scale bar: 10 μm. See also Supplementary Movie 1. **b**, Histograms of signal intensity and signal-to-background ratio of SunRISE labeled single mRNAs quantified with dNEMO^19^. Signal intensity is defined as the average of background-corrected pixel values within the area of each detected spot. Signal-to-background ratio is calculated as the ratio of average pixel intensity within an mRNA spot divided by the average intensity of background pixels in an annular ring surrounding the spot. (n=23 for cell numbers and n=5611 for spots numbers) **c**, Time course of single mRNA intensities for 8hrs time-lapse image. Red line indicates the median value of intensities for mRNA molecules in each frame and grey lines mark 25% and 75% quantiles of the intensities. **d**, HeLa cells transfected with MS2 (top) and MS2V6 (bottom) stem loops variants of SunRISE show similar characteristics and intensity distributions, quantified in **e**. See also Supplementary Figs. 2-6.

Puromycin is a translation inhibitor that causes premature termination of polypeptide chains, release of untranslated mRNAs, and induces formation of membrane-less cytosolic bodies called stress granules (SGs)^14, 15^. Recent studies have shown that G3BP protein modulates assembly of mRNAs and RNA-binding proteins into SGs^16-18^. *In vitro* reconstitution experiments have demonstrated that unfolded mRNAs can facilitate G3BP clustering and SG condensation^17^. However, it is difficult to assess the behavior of individual mRNAs during SG assembly *in vivo*. We therefore sought to observe SunRISE-labeled mRNAs and SG dynamics in the same cells exposed to puromycin-induced translational stress. Before stimulation with puromycin, single mRNA spots were observed and G3BP-mCh was diffuse throughout the cytoplasm. In response to puromycin, mRNA and G3BP condensed into droplets that form at different times between cells. The formation of mRNA droplets was independent of G3BP overexpression and did not co-localize with P-bodies (Supplementary Fig. 8). Although mRNA and G3BP droplets occasionally overlap, droplets were more often peripheral or within close vicinity to each other. Although these observations do not preclude roles for other mRNAs in SG assembly, our results show that mRNA condensates can be distinct from SG droplets and P-bodies (Fig. 3a and Supplementary Movie 2).

**Fig. 3:**
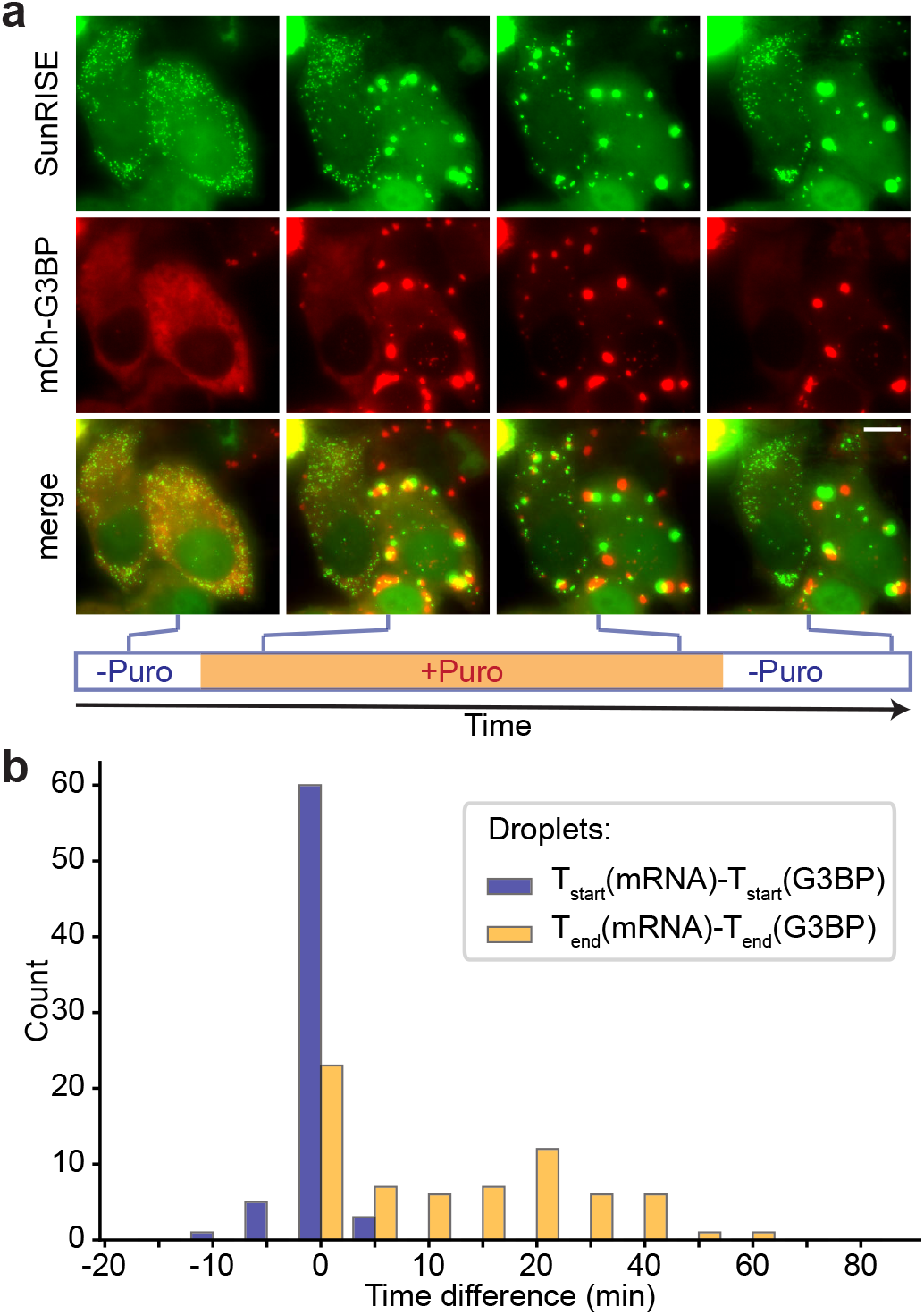
Long-term dynamics of stress granules and mRNA condensates following translational inhibition. **a**, Maximum intensity projection images of HeLa cells transfected for 24 hours with optimal SunRISE components and G3BP-mCherry to visualize stress granules. Cells were stimulated with puro (10 μg/ml) for 3hrs and then recovered in fresh media for 3 hours to observe the formation and dissolution of SG and mRNA droplets (See also Supplementary Movie. 2). Scale bar: 10 μm. **b**, Histograms of difference in timing of formation and dissolution of mRNA and G3BP droplets in single HeLa cells treated as described in panel a. See also Supplementary Figs. 9 and 10.

After puromycin removal, many stress granules disassemble and mRNA droplets dissipate back into single mRNAs (Fig. 3a). To establish differences between mRNA droplets and SGs, we exposed cells to a pulse of puromycin and measured the time of droplet formation and dissipation in each cell. Although mRNA and SG droplets form at the same time in most cells, SGs tend to dissipate earlier than mRNA droplets in cells that recover from stimulation (Fig. 3b and Supplementary Fig. 9). Furthermore, the duration of G3BP droplets is a strong predictor of whether a cell recovers from puromycin-induced condensates. In timescales of our experiments, when SGs persist for longer than 150 minutes in a cell, G3BP and mRNA droplets are unlikely to dissipate (Supplementary Fig. 10). Together, these results demonstrate that puromycin-induced mRNA and G3BP droplets are reversible and capable of independent phase separations.

Here, we described a live-cell reporter system for imaging mRNA over long periods of time. Through computational and experimental optimization, the reporter enables unambiguous detection of single mRNA molecules and does not extend mRNA perturbations beyond well-established 24x stem loop arrays. An advantage of SunRISE is that it is compatible with standard epifluorescence microscopy conditions used for live-cell experiments and therefore broadly accessible. We expect that SunRISE will enable dynamical single-cell studies of mRNA regulation and transport in addition to phase separation events with applications across biological disciplines.

## Supporting information

Integrated supplementary information

## Acknowledgements

We thank members of the Lee lab, Dr. Yi-Jiun Chen, Dr. Xiao-lun Wu, and Dr. Qiuhong Zhang for many helpful discussions. This work was supported by generous funding to RECL from NIH grant R35-GM119462 and the Alfred P. Sloan Foundation.

## Notes

### Competing Interest Statement

The authors have declared no competing interest.

