## Supplementary material for "SunRISE: long-term imaging of individual mRNA molecules in living cells": Integrated supplementary information

Supplementary figures and text:

| | |
|---|---|
| **Supplementary Figure 1** | Simulations to identify expression ratios of SunRISE components for consistent and high signal intensity. |
| **Supplementary Figure 2** | Varying promoters for scAB-GFP and 10xSunTag-PCP corroborates model predictions. |
| **Supplementary Figure 3** | The preliminary version of SunRISE labels only cytoplasmic mRNA because SunTag-PCP is excluded from nuclei. |
| **Supplementary Figure 4** | Modifications to SunTag-PCP alter sub-cellular distribution of SunRISE components. |
| **Supplementary Figure 5** | Optimized SunRISE detects mRNA in both cytoplasm and nucleus, verified by smFISH. |
| **Supplementary Figure 6** | Model calibrated to MS2V6-MCP affinity is comparable with previous observations for PP7. |
| **Supplementary Figure 7** | MoonRISE, MoonTag variant of SunRISE, provides an orthogonal tool for labeling of single mRNA molecules. |
| **Supplementary Figure 8** | SunRISE condensates require mRNA, are independent of G3BP overexpression, and do not co-localize with P-bodies. |
| **Supplementary Figure 9** | Difference in timing for formation and dissolution of mRNA and G3BP droplets in biological replicates. |
| **Supplementary Figure 10** | Duration of G3BP droplets predicts whether cells recover from puromycin stimulation. |
| **Supplementary Table 1** | Summary of SunTag/PCP plasmid variants. |
| **Supplementary Movie Legends** | |

#### Suplementary figure 1

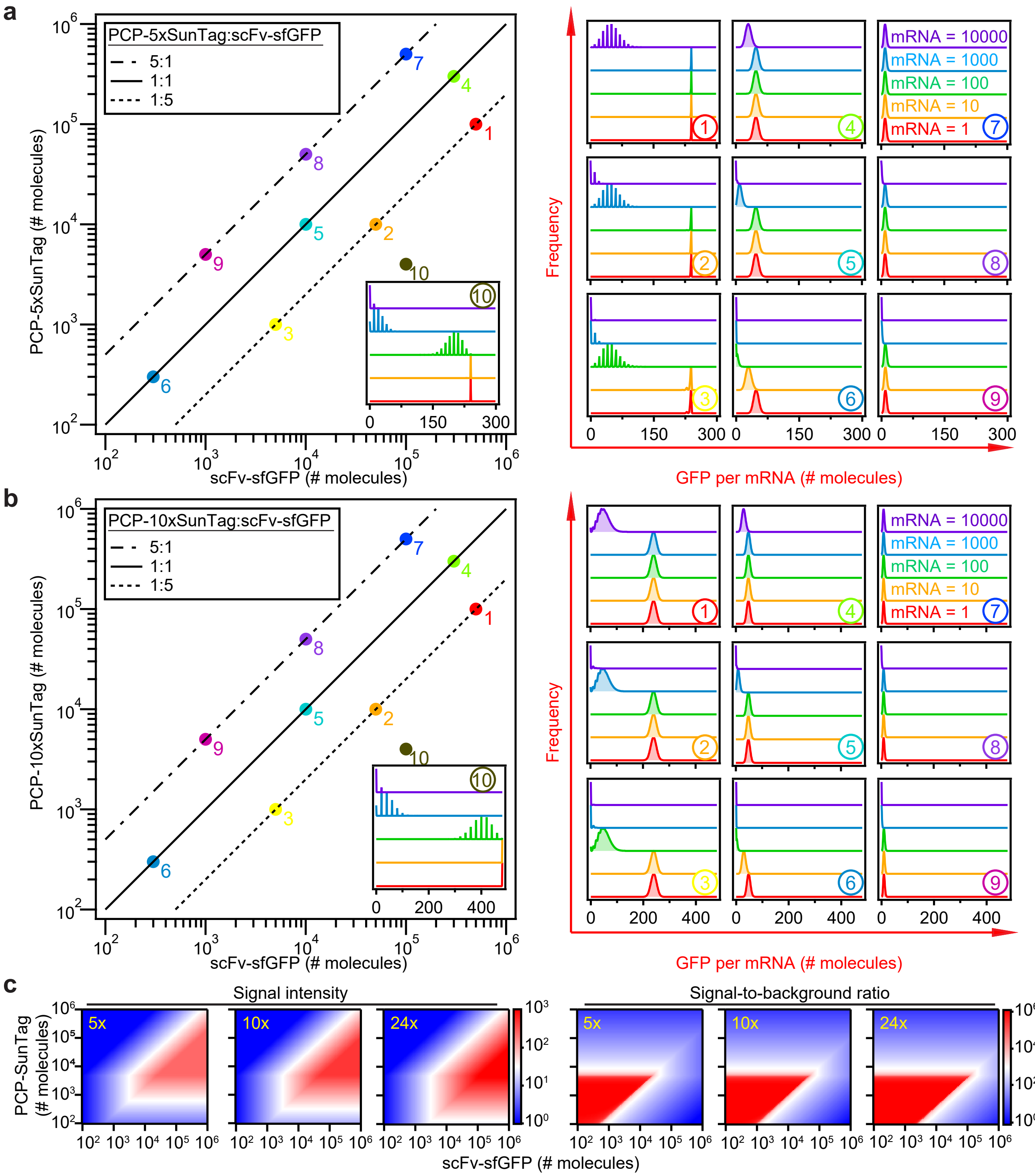

**Supplementary Figure 1. Simulations to identify expression ratios of SunRISE components for consistent and high signal intensity.**

**a and b,** Parameter sweeps in the space spanned by number of molecules for scFV-GFP and ‘n’xSunTag-PCP (left) using a computational model to calculate the number of GFP molecules per mRNA. Parameter combinations 1-10 were selected to represent different ratios between scFv-GFP and ‘n’xSunTag-PCP (solid line 1:1, dotted line 5:1 and dash-dotted line 1:5) in plots of frequency versus intensity of GFP labeling (right). Different concentrations of mRNAs per cell are distinguished by different colors. (a, 5xSunTag; b, 10xSunTag; See also, Fig. 1 for 24xSunTag). **c,** Heatmaps of mean number of GFP molecules per mRNA (left) and signal-to-background ratio (right) in the parameter space spanned by the number of molecules for scFv-GFP and ‘n’xSunTag-PCP (from left to right, 5x, 10x and 24x). Signal-to-background ratio is calculated as ratio of scFV-sfGFP molecules bound to an mRNA divided by the average intensity of unbound scFV-sfGFP in the background. Heatmaps were calculated assuming 100 mRNA molecules per cell.

#### Suplementary figure 2

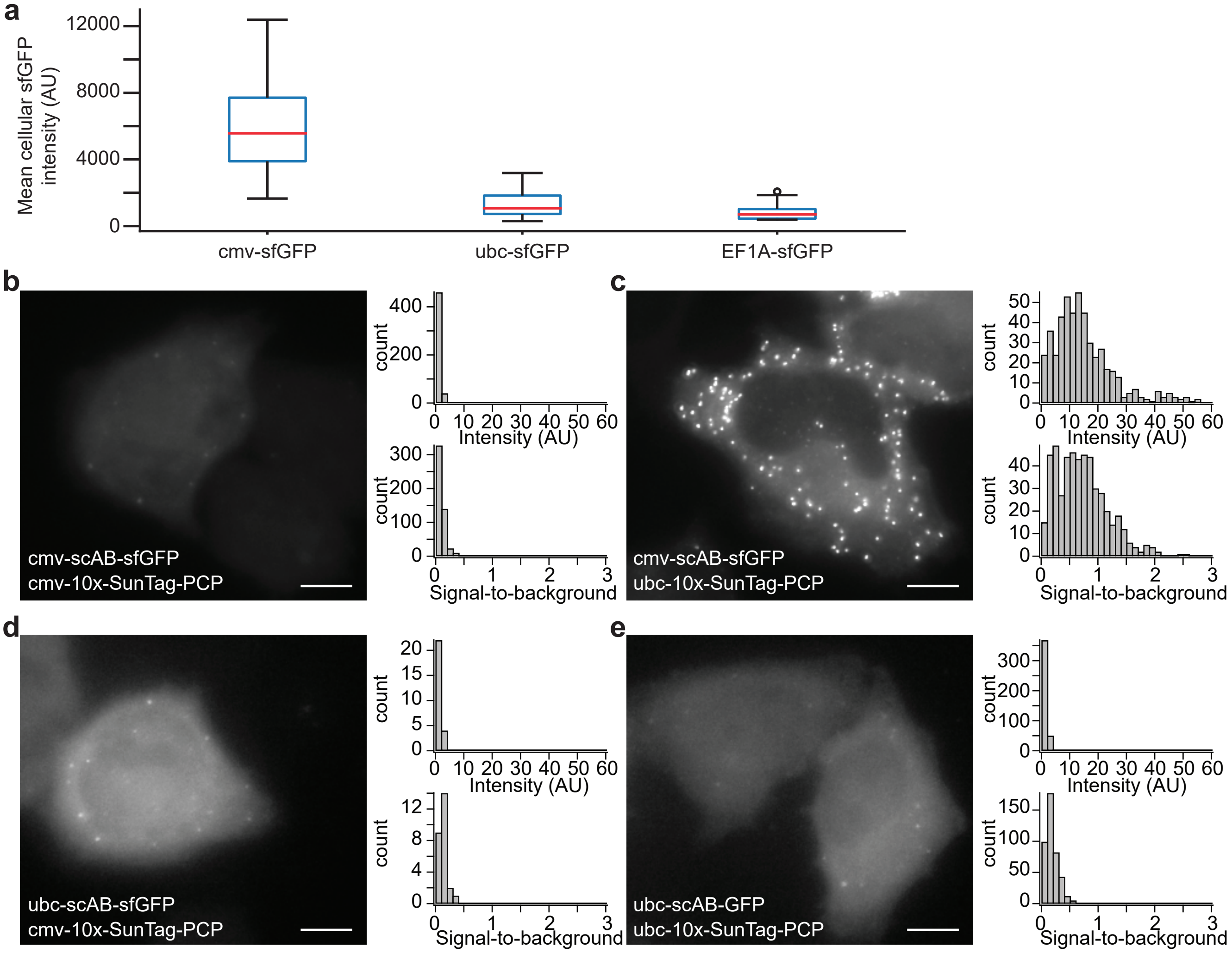

**Supplementary Figure 2. Varying promoters for scAB-GFP and 10xSunTag-PCP corroborates model predictions.**

**a,** HeLa cells transfected with cmv-scAB-GFP, ubc-scAB-GFP, or EF1A-scAB-GFP were imaged 24 hours after transfection and mean GFP fluorescence intensity was measured in single cells. Boxplots (median and interquartile ranges) show expression variability for indicated promoters. The cmv promoter is expressed approximately 5-times and 7-times that of ubc and EF1A promoters respectively. **b-e,** HeLa cells transfected with phage-cmv-cfp-24xpp7 and indicated constructs were imaged by 60x wide-field microscopy 24 hours after transfection and imaged with identical settings, representative maximum intensity projections are shown. Images were analyzed with dNEMO to quantify mRNA spots. Signal intensity is defined as the average of background-corrected pixel values within the area of each detected spot. Signal-to-background ratio is calculated as ratio of signal intensity divided by the average intensity of background pixels in an annular ring surrounding the spot. Note that images in (d) and (e) were contrast-enhanced for visualization. The combination of cmv-scAB-GFP and ubc-10xSunTag-PCP enabled robust detection of diffraction-limited mRNA spots and shows the highest signal intensity and signal-to-background ratio. Scale bar: 10 μm.

### Suplementary figure 3

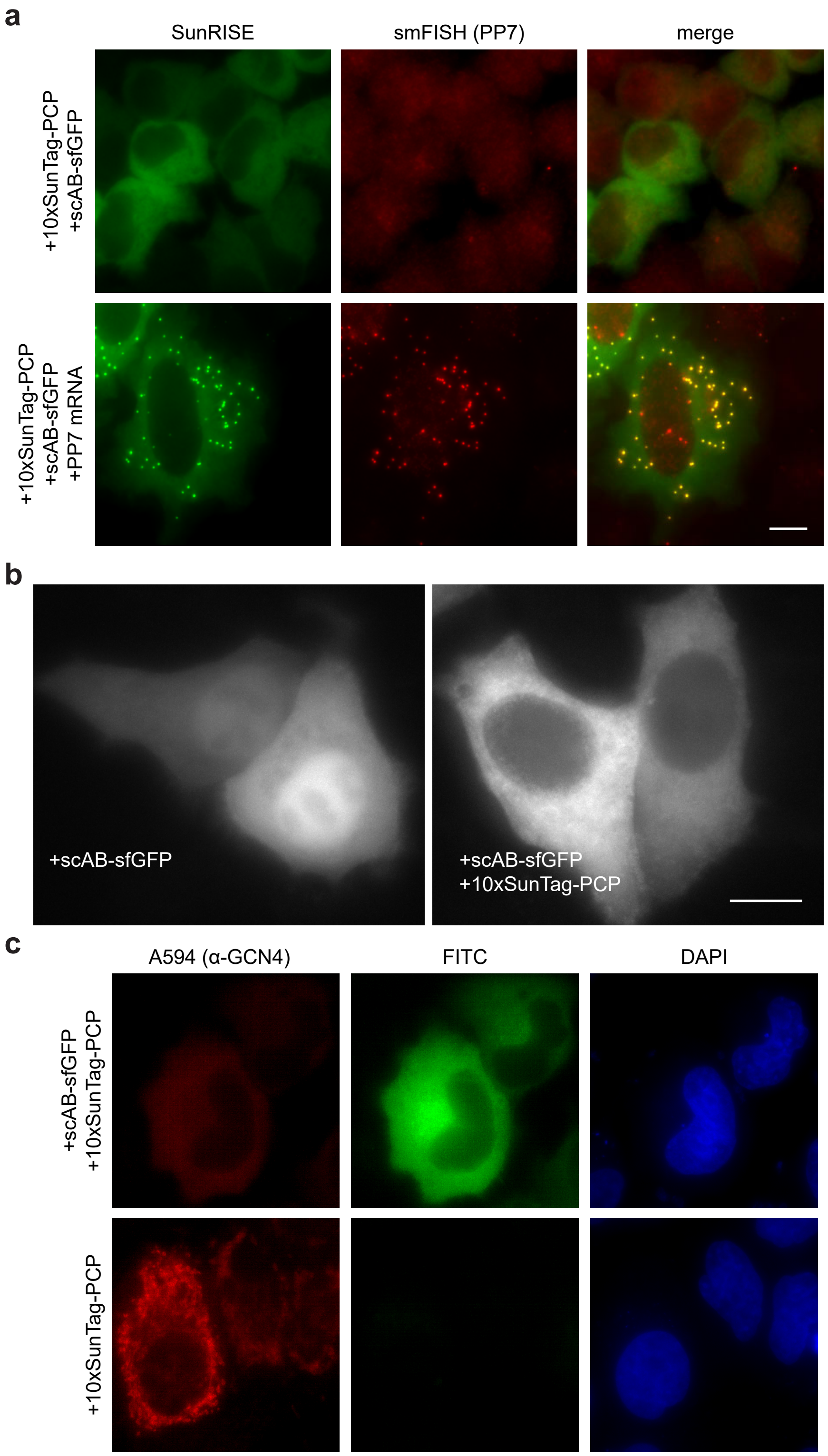

**Supplementary Figure 3. The preliminary version of SunRISE labels only cytoplasmic mRNA because SunTag-PCP is excluded from nuclei.**

**a,** Maximum intensity projection images of smFISH performed with probes against the pp7 stem-loops on HeLa cells transfected with indicated constructs are shown. The combination of cmv-scAB-GFP and ubc-10xSunTag-PCP allows visualization of cytoplasmic mRNA, but not mRNAs in nucleus. From left to right, FITC, A594, and merged channels are shown. **b,** HeLa cells were transfected with cmv-scAB-GFP only (left) or co-transfected with a combination of cmv-scAB-GFP and ubc-10xSunTag-PCP (right) and imaged after 24 hours. Co-expression of scAB-GFP and SunTag-PCP depletes scAB-GFP from the nucleus through interaction with SunTag-PCP. **c,** HeLa cells transfected with cmv-scAB-GFP and ubc-10xSunTag-PCP (top) and ubc-10xSunTag-PCP only (bottom) were stained for SunTag using a α-GCN4 antibody. Since scAB-sfGFP competes with the α-GCN4 antibody for the GCN4 epitope, co-expressing cells are lower intensity in the A594 channel (top left). In both conditions, PCP-10xSunTag is predominantly cytoplasmic regardless of whether scAB-GFP is expressed in the same cell. Scale bars: 10 μm for all.

**Suplementary figure 4**

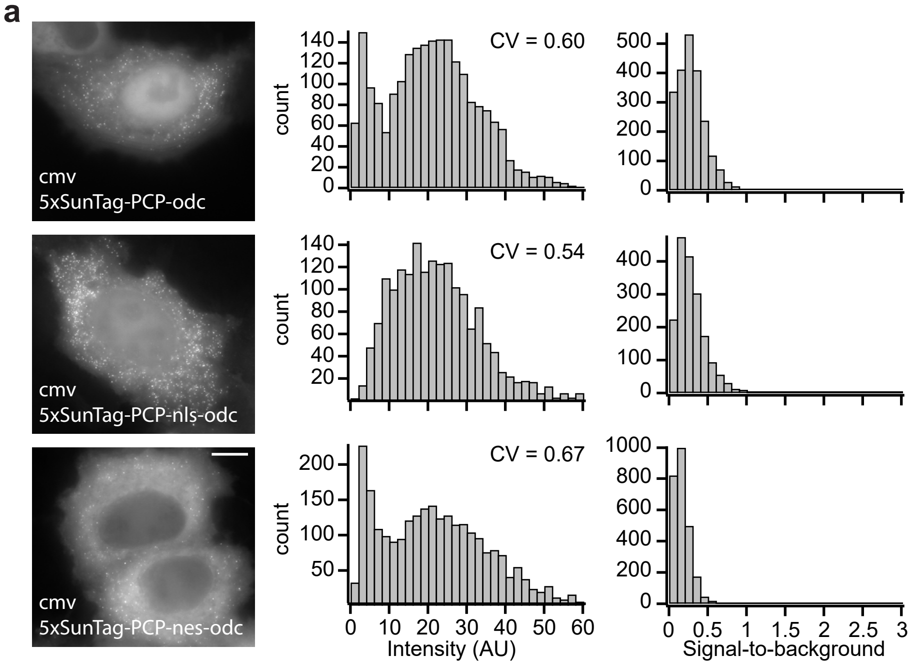


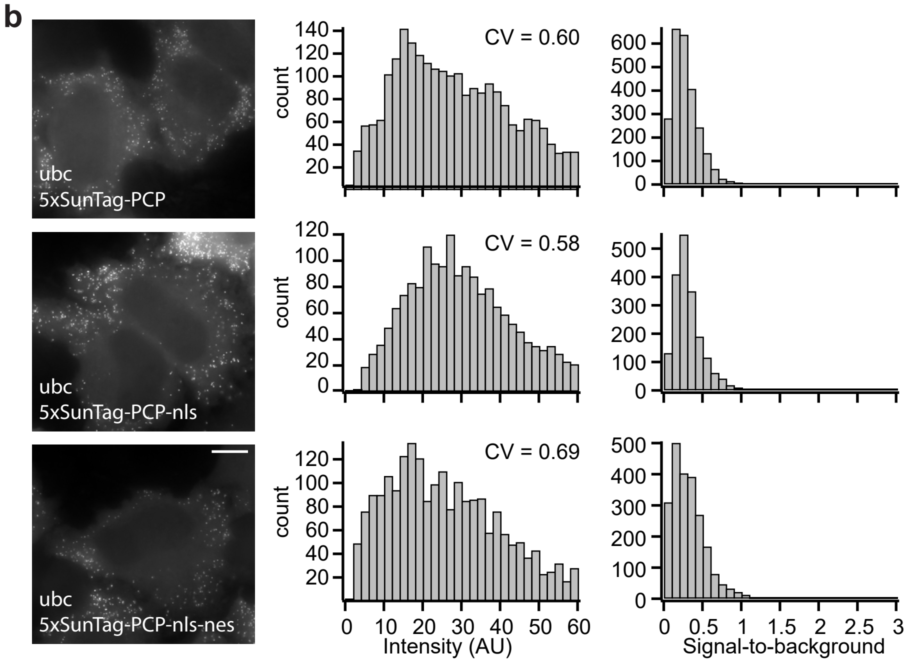

**Supplementary Figure 4. Modifications to SunTag-PCP alter sub-cellular distribution of SunRISE components.**

**a-b,** HeLa cells transfected with cmv-scAB-GFP, phage-cmv-cfp-24xpp7 and indicated constructs were imaged with identical settings by 60x wide-field microscopy 24 hours after transfection and quantified with dNEMO. Maximum intensity projections of representative cells are shown. Signal intensity is defined as the average of background-corrected pixel values within the area of each detected spot. Signal-to-background ratio is calculated as ratio of signal intensity divided by the average intensity of background pixels in an annular ring surrounding the spot. Coefficient of variation (CV) is the ratio of the standard deviation to the mean. ODC acts as a degron to reduce the expression level of SunTag-PCP driven by cmv promoter; nls, nuclear localization signal; nes, nuclear export signal. Scale bar: 10 μm. See also Supplementary Table 1.

**Suplementary figure 5**

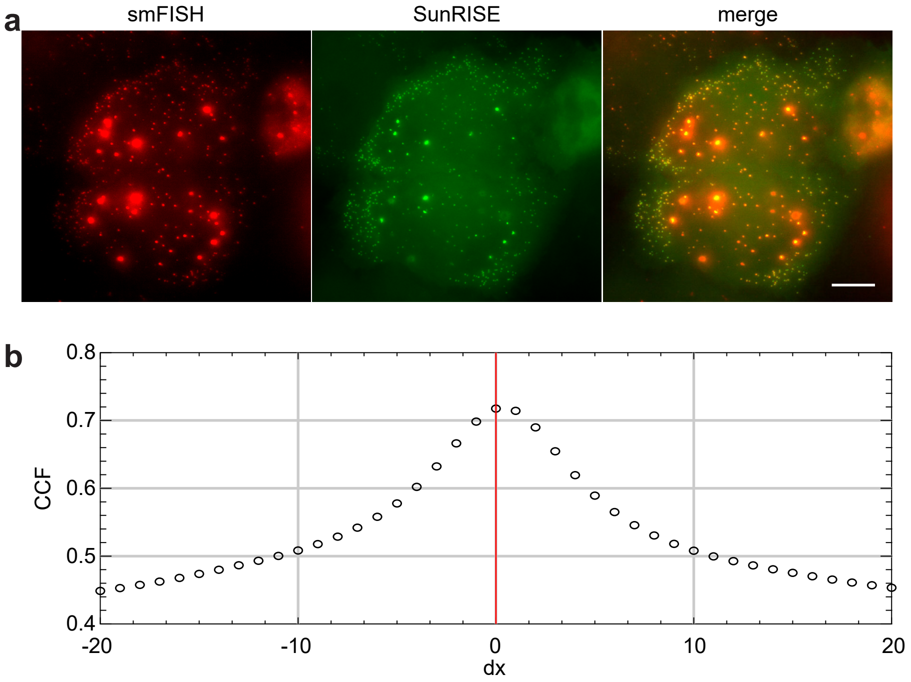

**Supplementary Figure 5. Optimized SunRISE detects mRNA in both cytoplasm and nucleus, verified by smFISH.**

**a,** Maximum intensity projection images of smFISH performed with probes against the pp7 stem-loops on HeLa cells transfected with phage-cmv-cfp-24xpp7, ubc-nls-PCP-5xSunTag and cmv-scAB-GFP (SunRISE) are shown. From left to right, FITC channel, A594 channel and merged channel. Scale bar: 10 μm. **b,** Cross-correlation functions (CCFs) calculated using Van Steensel's approach in the JACoP ImageJ plugin[1]. X-axis variable, dx, defines the pixel shift distance between two fluorescence channels and red line marks dx = 0. Black circles are Pearson's coefficients calculated for each pixel shift. The distribution of Pearson correlations peak near dx = 0, showing almost complete co-localization between smFISH and SunRISE channels. The peak value of the distribution is less than 1, indicating differences in fluorescence intensities between the two fluorescence channels.

#### Suplementary figure 6

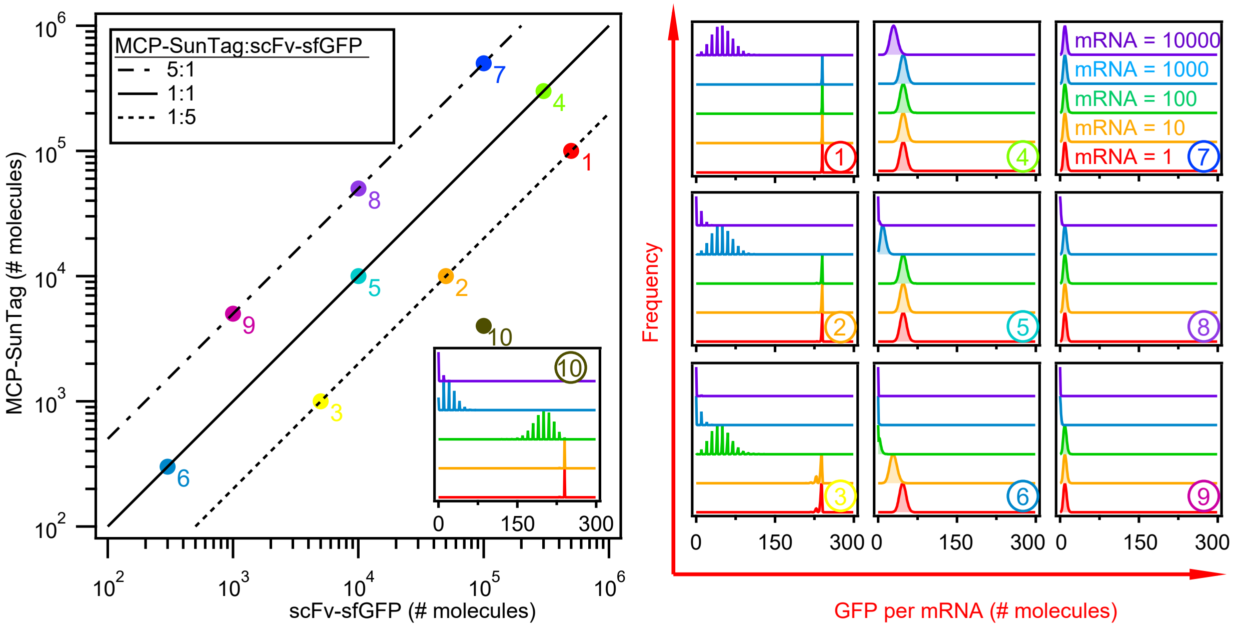

**Supplementary Figure 6. Model calibrated to MS2V6-MCP affinity is comparable with previous observations for PP7.**

Parameter sweeps in the space spanned by the number of molecules for scFV-GFP and 5xSunTag-MCP (left) using a computational model to calculate number of GFP molecules per mRNA. Parameter combinations 1-10 were selected to represent different ratios between scFV-GFP and 5xSunTag-MCP (solid line 1:1, dotted line 5:1 and dash-dotted line 1:5) and different expression levels and frequency plots are shown (right). Different concentrations of mRNAs per cell are distinguished by different colors.

#### Suplementary figure 7

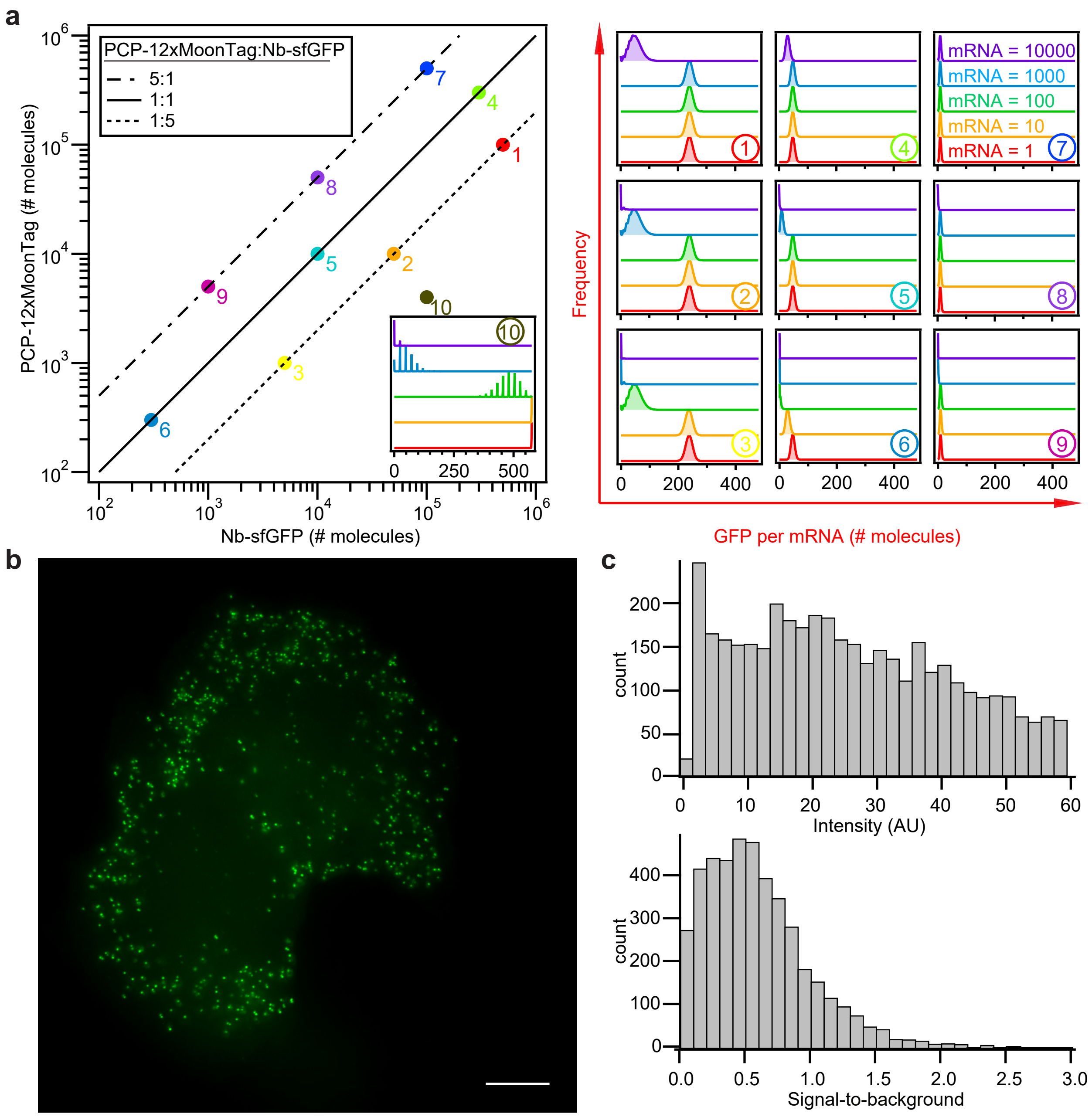

**Supplementary Figure 7. MoonRISE, MoonTag variant of SunRISE, provides an orthogonal tool for labeling of single mRNA molecules.**

**a,** Parameter sweeps in the space spanned by the number of molecules for Nb-gp41-GFP and 12xMoonTag-PCP (left) using a computational model to calculate number of GFP molecules per mRNA. Parameter combinations 1-10 were selected to represent different ratios between Nb-gp41-GFP and 12xMoonTag-PCP (solid line 1:1, dotted line 5:1 and dash-dotted line 1:5) in plots of frequency versus intensity of GFP labeling (right). Different concentrations of mRNAs per cell are distinguished by different colors. **b,** HeLa cells transfected with cmv-Nb-gp41-GFP, phage-cmv-cfp-24xpp7 and ubc-nls-PCP-12xMoonTag were imaged with 60x wide-field microscope 24 hours after transfection and quantified with dNEMO in **c,**. Scale bar: 10 μm.

#### Suplementary figure 8

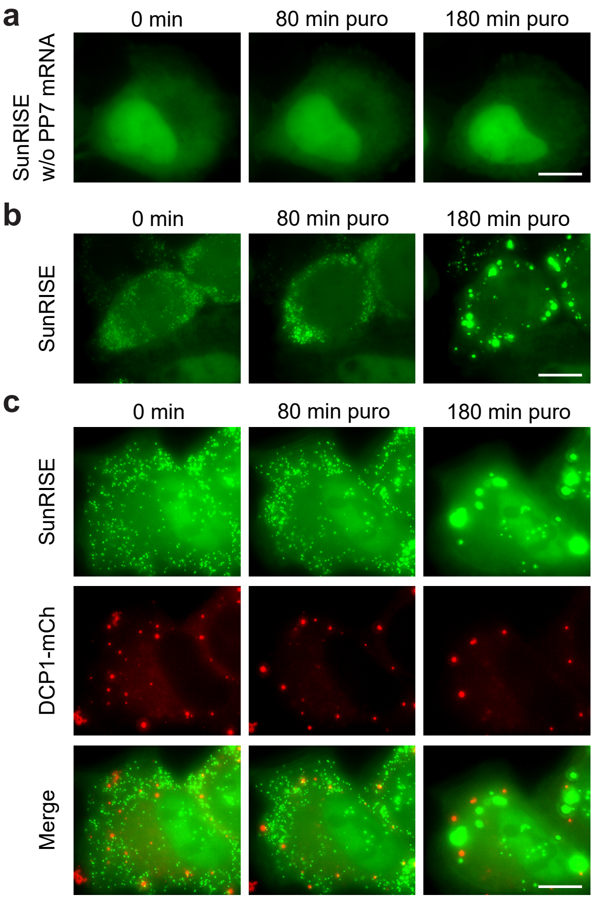

**Supplementary Figure 8. SunRISE condensates require mRNA, are independent of G3BP overexpression, and do not co-localize with P-bodies.**

Representative maximum intensity projection images of HeLa cells transfected for 24 hours with indicated constructs. For all experiments, cells were stimulated with puromycin (10 μg/ml) for 3hrs and then recovered in fresh media for 3hrs to observe the formation and dissolution of mRNA droplets. **a,** Cells with cmv-scAB-GFP and ubc-nls-PCP-5xSunTag, but not expressing PP7 stem loops show no condensates in response to puromycin. **b,** Cells with all SunRISE components that do not overexpress G3BP still form condensates similar to cells that do overexpress G3BP (see Fig. 3). **c,** Cells expressing SunRISE components and Dcp1-mCherry to mark P-bodies do not form new P-bodies following puromycin stimulation. SunRISE condensates and pre-existing P-bodies do not co-localize. Scale bar: 10 μm.

#### Suplementary figure 9

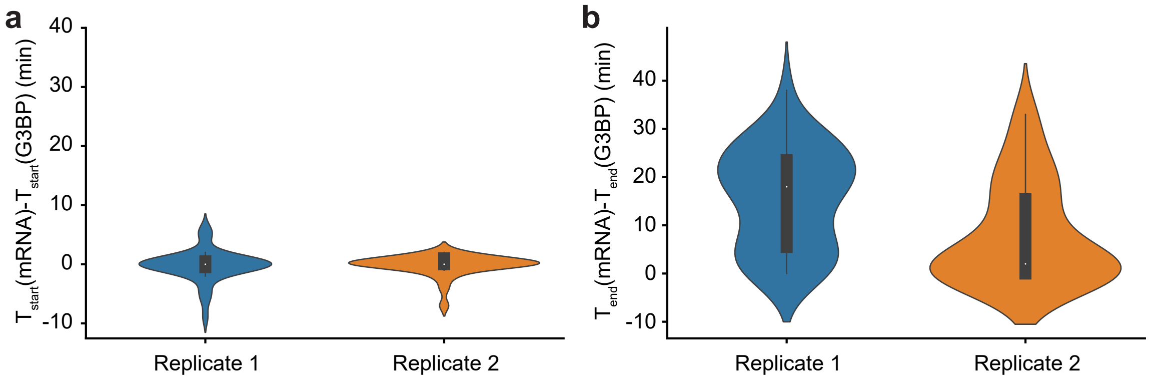

**Supplementary Figure 9. Difference in timing for formation and dissolution of mRNA and G3BP droplets in biological replicates.**

Violin plots of differences between starting time ($T_{start}$) (**a,**) and time of dissipation ($T_{end}$) (**b,**) for mRNA and G3BP droplets observed in the same cell. HeLa cells were treated with a 3-hour pulse of puromycin and allowed to recover for 3 hours post stimulation.

#### Suplementary figure 10

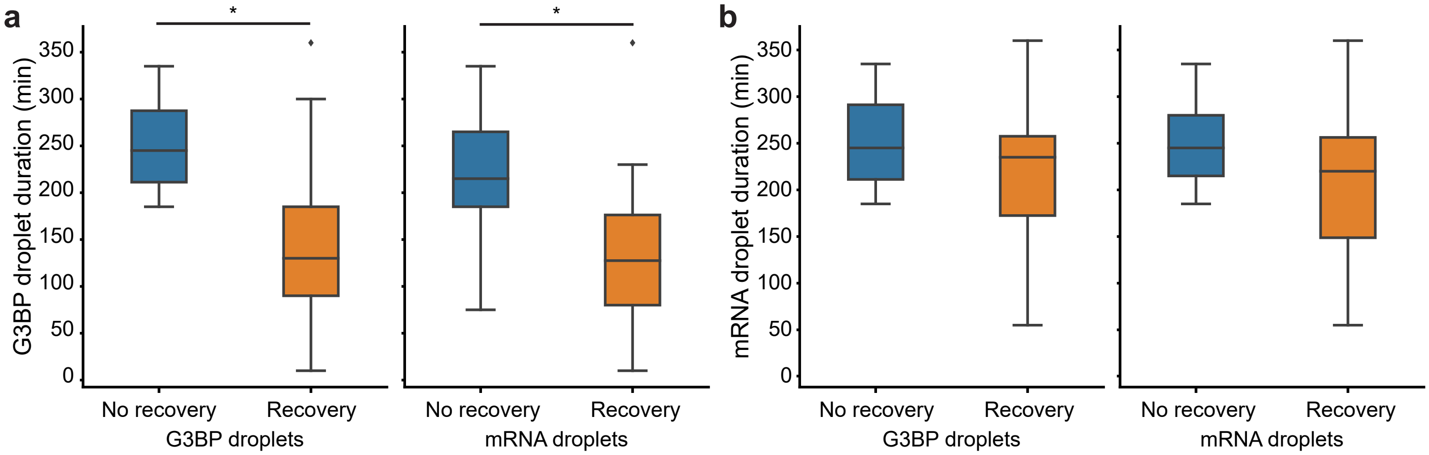

**Supplementary Figure 10. Duration of G3BP droplets predicts whether cells recover from puromycin stimulation.**

HeLa cells were treated with a 3-hour pulse of puromycin and allowed to recover for 3 hours post stimulation. Boxplots (median and interquartile range) of droplet duration for **a,** G3BP and **b,** mRNA. Single cells were partitioned into two categories. The ‘recovery’ designation for the indicated reporter (G3BP or mRNA on the x-axis) identifies cells where G3BP droplets resolve or where single mRNA molecules were observed to dissipate from droplets within the duration of the experiment. * $p < 10^{-5}$, t test.

**Supplementary Table 1. Summary of SunTag/PCP plasmid variants.**

SunTag-PCP sequence modifications include varying promoters (ubc and cmv), nuclear localization sequences (nls), nuclear export sequences (nes) and ornithine decarboxylase (ODC). Note that “Limited” indicates that significant cell-to-cell variability or lower number than expected was observed for the associated property.

| PCP/SunTag variants | Nuclear GFP | Nuclear spots | Cytoplasmic spots |
|---|---|---|---|
| **ubc-PCP-10xSunTag-nls** | No | Limited | Yes |
| **ubc-PCP-10xSunTag-nls-nes** | No | Limited | Yes |
| **cmv-PCP-10xSunTag-ODC** | Yes | No | Yes |
| **cmv-PCP-10xSunTag-nls-ODC** | Yes | Yes | Yes |
| **cmv-PCP-10xSunTag-nes-ODC** | Yes | Limited | Yes |
| **cmv-PCP-10xSunTag-nls-nes-ODC** | Yes | Limited | Yes |
| **ubc-PCP-10xSunTag-ODC** | Limited | No | Limited |
| **ubc-PCP-10xSunTag-nls-ODC** | Yes | Limited | Limited |
| **ubc-PCP-10xSunTag-nes-ODC** | Limited | No | Limited |
| **ubc-PCP-10xSunTag-nls-nes-ODC** | Yes | No | Limited |
| **cmv-PCP-5xSunTag-ODC** | Limited | No | Yes |
| **cmv-PCP-5xSunTag-nls-ODC** | Yes | No | Yes |
| **cmv-PCP-5xSunTag-nes-ODC** | No | No | Yes |
| **ubc-5xSunTag-PCP-nls** | No | Limited | Yes |
| **ubc-5xSunTag-PCP-nls-nes** | No | Limited | Yes |
| **ubc-nls-5xSunTag-PCP-nls** | Yes | No | Yes |
| **ubc-nls-5xSunTag-PCP-2xnls** | Yes | No | Limited |
| **ubc-2xnls-5xSunTag-PCP-2xnls** | Yes | No | No |
| **ubc-PCP-5xSunTag** | No | No | Yes |
| **ubc-nls-PCP-5xSunTag** | Yes | Yes | Yes |
| **ubc-nls-PCP-5xSunTag-nes** | Yes | No | Yes |

#### Supplementary movie legends

**Supplementary Movie 1. The optimized SunRISE design enables long-term imaging of single mRNA molecules throughout the cell, related to Fig. 2.**

Movie of maximum intensity projection images for HeLa cells expressing SunRISE components. Cells were imaged with a 10-minute interval between frames.

**Supplementary Movie 2. HeLa cells expressing SunRISE and G3BP-mCherry show formation and dissolution of SG and mRNA droplets upon stimulation with a puromycin pulse, related to Fig. 3a.**

Movie of maximum intensity projection images for HeLa cells expressing SunRISE components (green) and mCherry-G3BP (red). Cells were imaged with a 5-minute interval between frames. For this experiment, puromycin was introduced at t = 0 minutes and removed at t = 270 minutes.
