## Supplementary material for "SunRISE: long-term imaging of individual mRNA molecules in living cells": Methods

#### Plasmids construction

The 24xSunTag-PCP plasmids were constructed with 24xGCN4 repeats flanked by HindIII and BamHI sites and coat protein flanked by BamHI and EcoRI sites in a pcDNA3 vector. Ubc promoter was PCR amplified from phage-ubc-nls-ha-pcp-gfp (Addgene #64539) and inserted between MluI and HindIII sites to make ubc-nxSunTag-PCP. 5xSunTag and 10xSunTag variants were generated by PCR amplification of 5xGCN4 and 10xGCN4 from pcDNA4TO-5xGCN4_v4-kif18b-24xPP7 (Addgene #74927) and pcDNA4TO-mito-mCherry-10xGCN4_v4 (Addgene #60914) and replacing 24xSunTag respectively. SV40NLS, a 57bp NES signal (ATGAACCTGGTGGACCTCCAAAAGAAGCTGGAGGAGCTGGAGCTGGACGAGCAG CAG) or NES from HIV Rev protein and ODC fragment amplified from pEF-24xV4-ODC-24xPP7 (a gift from Dr. Xiaowei Zhuang's lab) were added at the C-terminus of nxSunTag-PCP between EcoRI and XbaI for various modifications. The ubc-nls-PCP-5xSunTag plasmids were created by replacing gfp sequence in phage-ubc-nls-ha-pcp-gfp with 5xGCN4 flanked by BamHI and BsrGI sites. Similarly, ubc-nls-MCP-5xSunTag were obtained by insertion of 5xGCN4 to UbC-NLS-HA-MCP-YFP (Addgene #31230) after digestion of XbaI and BsrGI restriction enzymes.

cmv-sfGFP-GB1-scAB was assembled by inserting sfGFP-GB1 fragment from pHRdSV40-scFv-GCN4-sfGFP-VP64-GB1-NLS (Addgene #60904) and scAB fragment from phage UbiC scAB-GFP (Addgene #104998) into a pcDNA3 vector.

phage-cmv-cfp-24xms2 (Addgene #40651) and phage-cmv-cfp-24xpp7 (Addgene #40652) act as reporter mRNA labeled with different stem-loops. 24xMS2V6 stem-loops from pET259-pUC57-24xMS2V6 (Addgene #104391) was amplified using BamHI and SacII sites to label cfp the same way as other two stem-loops.

For MoonRISE variant, cmv-sfGFP-GB1-Nb-gp41 was created by replacing scAB with Nb-gp41 from Nb-gp41-Halo (MoonTag-Nb-Halo) (Addgene #128603). Ubc-nls-PCP-12xMoonTag is made by using 12xMoonTag-12xSunTag-kif18b-24xPP7 (Addgene #128606) as backbone and removing sequences after 12xMoonTag and inserting stop codon and inserting ubc-nls-PCP fragment between SpeI and HindIII sites in front of 12xMoonTag.

G3BP1 and Dcp1 amplified from pEGFP-C1-G3BP1-WT (Addgene #135997) and pT7-EGFP-C1-HsDCP1a (Addgene #25030) were subcloned into the H2B-mCherry (Addgene #20972) using HindIII/BamHI sites and KpnI/AgeI sites respectively.

**Cell culture and puromycin treatment**

HeLa cells were maintained in DMEM supplemented with 10% fetal bovine serum, 1% streptomycin/penicillin and 1% L-glutamine at 37℃ with 5% humidified $CO_2$. Cells were plated on 96-well glass bottom plates (Matriplate) for fixed-cell and live cell imaging experiments. Puromycin (Gibco) was diluted into DMEM and 30 μL of 100 μg/ml mixture was added into wells containing 270 μL of growth media to yield final concentration of 10 μg/ml. Puromycin containing media were replaced with pre-warmed DMEM media to allow cells to recover from translational stress.

### Live cell imaging and quantification of single mRNA spots with dNEMO

HeLa cells were plated on 96-well glass bottom plates (Matriplate) at the density of $8x10^3$~ $1x10^4$ per well. Transient transfection was performed with Fugene HD (Promega) 24hrs later according to manufacturer's protocol. A mixture of plasmids comprising SunTag system with equal amount for each was first made in Opti-MEM then Fugene HD was added and incubated for 15 mins at room temperature. The total amount of DNA and Fugene HD can be optimized accordingly. 24hrs after transfection cells were imaged using a DeltaVision Elite microscope with a 60x objective (1.42 NA; Olympus) and temperature-matched oil in an environmentally controlled chamber (37°C, 5% $CO_2$). Z-stacks of 5 images with 1 μm interval were acquired for quantification with dNEMO. Cell segmentation was manually performed in dNEMO and the spot detection parameters are set as default. For quantification in systematic optimization of 5xSunTag-PCP, on average 17 cells and at least 1800 spots in each condition were used for plotting histograms. For comparison of promoters, mean fluorescence intensity was measured using ImageJ for a fixed region in cytoplasmic areas for each cell.

### Quantification of timing for droplets formation and dissolution

The quantification was performed on the pool of cells that survive the puromycin pulse treatment till the end of 6-hr movie and at least in one channel the droplets formation and dissolution were observed. The time of start in G3BP channel was defined as the frame in which fluorescent puncta form and the time of end as the frame in which G3BP becomes diffuse again. The time of start in SunRISE channel was defined as the frame in which the size of spot grows and the shape becomes irregular and diffraction-limited spots are not resolvable and the time of end as the frame in which single mRNA spots reemerge. Note that till the end of movies, some droplets in SunRISE channel are still in the process of dissolution, which means we observe the size of droplet reducing and mRNA releasing from droplets. We use the last frame as the time of end for this scenario.

**smFISH probes and image acquisition**

Five 3’ Cy5 fluorescently labeled DNA oligos[1](ggcaattaggtaccttagg, catatcgtctgctcctttc, gagtcgacctgcagggag, atatgctctgctggtttc, atactgcagccagcgagc) as smFISH probes against PP7 stem-loops were synthesized by Genewiz. HeLa cells were plated on 96-well glass bottom plates and transfected with SunRISE components for 24hrs. Cells were then fixed with 3.7% formaldehyde, washed three times in 1xPBS for 5 mins each and permeabilized in 70% (v/v) EtOH overnight at 4℃. The hybridization was then performed overnight at 37℃ with 100 nM probes in 2XSSC with 10% formamide and 10% dextran sulfate. Nuclei were labeled with 5 ng/mL DAPI in wash buffer during the wash step after the hybridization. Cells were finally imaged in Glox buffer[2] using a 60X objective on a DeltaVision microscope. Z-stack images of both FITC channel (SunRISE) and Cy5 channel (smFISH) were collected.

**Fixed-cell Immunofluorescence**

HeLa cells were plated on 96-well glass bottom plates and transfected with 10xSunTag-PCP or the combination of scAB-GFP and 10xSunTag-PCP for 24hrs. Cells were then fixed with 3.7% formaldehyde for 10 minutes, rinsed three times in 1xPBS for 5 mins each and incubated in 100% methanol for 10 min. Cells were washed three times in PBST (1XPBS 0.1% Tween 20) for 5 mins each and then a primary antibody α-GCN4 (AbsoluteAntibody C11L34) diluted 1:100 in 3% BSA PBST was applied and incubated overnight at 4°C. After three washes with PBST, cells were incubated with secondary antibody (3% BSA PBST with 1:1000 Alexa594-conjugated anti-mouse IgG antibody) for one hour at room temperature. Nuclei were counterstained with 1 µg/mL Hoechst in PBST during the wash step after secondary antibody. Cells were finally imaged using a 60X objective on a DeltaVision microscope.

**Model-based calculation of signal intensity and signal-to-background values**

In the calculation, we assume that all dynamic processes, e.g. the expression of proteins, the transcription of mRNAs and the binding/unbinding of proteins, have reached equilibrium. The dissociation constant of scFv and GCN4, can be written as:

$$\kappa_{D1} = \frac{[scFv] \cdot [GCN4]}{[scFv \cdot GCN4]} = \frac{(c_1 - p_1 N_1 c_2) \cdot (1 - p_1) N_1 c_2}{p_1 N_1 c_2}$$

where $c_1$ is the molar concentration scFv. $c_2$ is the molar concentration of PCP-SunTag. $N_1$ is the number of GCN4 peptides on the PCP-SunTag. $p_1$ is the probability of a GCN4 binding site occupied by the scFv. Thus, we obtain:

$$p_1 = \frac{1}{2N_1 c_2}\left(\kappa_{D1} + N_1 c_2 + c_1 - \sqrt{(\kappa_{D1} + N_1 c_2 + c_1)^2 - 4c_1 N_1 c_2}\right)$$

Similarly, the probability of a PP7 binding site occupied by a PCP is:

$$p_2 = \frac{1}{2N_2 c_3}\left(\kappa_{D2} + N_2 c_3 + c_2 - \sqrt{(\kappa_{D2} + N_2 c_3 + c_2)^2 - 4c_2 N_2 c_3}\right)$$

Where $c_3$ is the molar concentration of mRNA, $N_2$ is the number of PP7 stem-loops on the mRNA, $\kappa_{D2}$ is the dissociation constant between PCP and PP7.

The number of scFv binding to a single PCP-SunTag satisfies a binomial distribution

$$B_1(n) = \binom{N_1}{n} p_1^n (1 - p_1)^{N_1 - n}$$

Similarly, the number of PCP-SunTag binding to a single mRNA molecule satisfies:

$$B_2(m) = \binom{N_2}{m} p_2^m (1 - p_2)^{N_2 - m}$$

The average number of scFv-GFP on a single mRNA molecule is thus

$$\bar{n} = \sum_{m=0}^{N_2}\left(m \cdot B_2(m) \cdot \sum_{n=0}^{N_1} n \cdot B_1(n)\right) = N_1 p_1 N_2 p_2$$

To calculate the distribution of number of scFv-GFP binding to a single mRNA molecule, we first calculate the probability of n scFv-GFP binding to m PCP-SunTag, which is

$$B_1^m(n) = \binom{mN_1}{n} p_1^n (1 - p_1)^{mN_1 - n}$$

Then the probability of n scFv-GFP on a single mRNA molecule is the sum of $B_1^m(n)$ of all possible m, which is:

$$P(n) = \sum_{m=0}^{N_2} B_2(m) \cdot B_1^m(n) = \sum_{m=0}^{N_2} \binom{N_2}{m} p_2^m (1-p_2)^{N_2-m} \cdot \binom{mN_1}{n} p_1^n (1-p_1)^{mN_1-n}$$

$$= (1-p_2)^{N_2} \left(\frac{p_1}{1-p_1}\right)^n \sum_{m=0}^{N_2} \binom{N_2}{m} \binom{mN_1}{n} \left(\frac{p_2}{1-p_2}\right)^m (1-p_1)^{mN_1}$$

The dissociation constants for scFv/GCN4 and PCP/PP7 are 0.38 nM[3] and 1 nM[4], respectively. The dissociation constants for Nb-gp41/MoonTag and MCP/MS2V6 are 30nM[5] and 2.4nM[6].

Considering that the signal from one mRNA molecule will spread to an ellipsoidal area, of which the size is determined by the Rayleigh radius, so we calculate the fluorescence intensity of this area in the presence and without the presence of a single mRNA molecule to calculate signal to background values. Here, the volume we chose for calculation is V=200 nm ×200 nm ×500 nm. If there is no mRNA in the area, the intensity is defined as the background intensity, $I_B$. Assuming that the intensity of single GFP-scFv is 1, the background intensity is the sum of free scFv-GFP molecules and the GFP-SunTag-PCP complex

$$I_B = N_A V(c_1 - N_1 p_1 c_2) + N_A V(c_2 - N_2 p_2 c_3) N_1 p_1$$

where $N_A$ is Avogadro constant.

With the intensity of an mRNA molecule as $I_{RNA} = N_1 p_1 N_2 p_2$ the signal-to-background is

$$\frac{I_{RNA}}{I_B} = \frac{N_1 p_1 N_2 p_2}{N_A V(c_1 - N_1 p_1 c_2) + N_A V(c_2 - N_2 p_2 c_3) N_1 p_1}$$

**Methods References**
